# Circadian Timekeeping in the Tropics: Rhythmic Transcriptome and Diurnal Regulatory Networks in *Rubroshorea leprosula*

**DOI:** 10.64898/2026.04.13.717981

**Authors:** Shashank Kumar Singh, Motohide Seki, Hiroshi Ito, Naoki Tani, Yuka Ikezaki, Kayoko Ohta, Suat Hui Yeoh, Ng Ching Ching, Akiko Satake

## Abstract

Circadian rhythms allow plants to regulate internal processes to align with daily rhythms in the environment. Comparing circadian clocks across environments is essential because latitude-dependent variation in light and temperature imposes distinct selection pressures that shape the evolution and function of circadian timing systems. However, **c**ircadian clock studies have largely focused on temperate and subtropical species, leaving transcriptional circadian networks in tropical plants under relatively constant environments poorly understood. We report the first comprehensive circadian transcriptome of the ecologically important dipterocarp species *Rubroshorea leprosula*, a dominant species in tropical rainforests in Southeast Asia. One sapling was sampled every four hours for 48 hours under constant darkness and used for transcriptomic analyses. Only 283 of 20,748 expressed genes (∼1.3%) exhibited significant circadian oscillations, with periods strongly concentrated between 23 and 25 h. Hierarchical clustering revealed four temporal clusters with alternate phases of expression and functional specialisation; morning clusters in processes related to light and chloroplast, midday clusters in hormone and signalling mechanisms, afternoon clusters in mitochondrial and peptide biosynthesis functions, and night clusters in protein quality control and autophagy. Comparative analysis identified clear orthologs for all major *Arabidopsis* circadian clock components (*LHY, CCA1, PRRs, TOC1, ELF4, LUX, GI, ZTL, RVE*s), with conserved synteny to *Parashorea chinensis*, a close relative of *R. leprosula*. Time-lagged cross-correlation (TLCC) network reconstruction identified a characteristic circadian topological similarity with Arabidopsis, including coupled morning and evening feedback loops and paralog expansion that maintained overall structure. Peak expression timing of these core clock genes in the tropical tree was largely consistent with that observed in *Arabidopsis thaliana*. In contrast to this conserved phase relationship, *Rubroshorea* orthologs exhibited reduced amplitudes and lower coefficients of variation in their circadian oscillations, suggesting diminished robustness of rhythmic gene expression. These findings demonstrate a conserved but regulated circadian mechanism in *R. leprosula*, in preparation for adaptation to tropical rainforest’s stable light and temperature regimes. This study lays the molecular foundation for circadian regulation in dipterocarps and offers a system for integrating rhythmic gene expression to ecological function and forest productivity in tropical communities.

## 1. Introduction

Circadian rhythms are endogenous, self-sustained oscillations that last approximately 24 hours, enabling organisms to anticipate and adapt to daily environmental fluctuations. The circadian clock coordinates plant’s key physiological and metabolic processes, including photosynthesis, hormone signalling, growth, and stress responses, optimizing fitness and productivity across diurnal cycles (Dodd et al., 2005; Inoue et al., 2017; Atamian et al., 2016; Greenham & McClung, 2015). Extensive studies in *Arabidopsis thaliana* have established the molecular framework of the plant circadian system, revealing complex transcriptional feedback loops and post-translational regulation that maintain rhythmic gene expression and phase synchronization with light-dark transitions (Webb et al., 2019; Nakamichi, 2020).

Recent theoretical work highlights that the adaptive value of circadian clocks is strongly shaped by environmental seasonality. In particular, the analysis of the evolutionary model of gene regulatory network demonstrated that self-sustained circadian oscillators are selectively favoured in environments with pronounced annual changes in photoperiod, where maintaining a stable internal phase becomes essential despite shifting sunrise and sunset times (Seki & Ito, 2022). Conversely, in aseasonal tropical regions with minimal variation in day length, damped or externally driven oscillators perform equally well, suggesting reduced pressure to maintain a robust endogenous rhythm. This raises an important evolutionary hypothesis: tropical plants may operate with a weaker or less autonomous circadian clock compared to temperate species, relying more heavily on direct environmental cues to regulate daily physiological processes.

Evidence from lower-latitude organisms further supports this evolutionary perspective. In aquatic plants such as duckweeds, circadian properties become increasingly irregular toward equatorial regions, indicating relaxed selection for precise endogenous timekeeping under weak seasonal variation (Isoda et al., 2022). Similarly, certain cyanobacterial lineages inhabiting tropical environments have evolutionarily lost circadian clock function altogether, suggesting that robust self-sustained oscillators are not universally required under stable environmental conditions (Holtzendorff et al., 2008; Flombaum et al., 2013). Together, these findings point to a general trend in which reduced seasonality is associated with attenuated or externally driven circadian dynamics across diverse taxa. Testing this hypothesis requires detailed time-series data from tropical taxa, which have remained largely unexplored to date.

However, detailed knowledge of circadian clock function is largely based on controlled environments and a few temperate model species, and understanding clock behavior in diverse, including tropical, natural environments remains limited (Laosuntisuk et al., 2023). Although limited in number, several studies in tropical plants indicate that circadian clocks remain functionally important even under weak seasonal variation. For example, genomic and expression analyses in the tropical tree Carica papaya revealed that core circadian clock genes (including *Late Elongated Hypocotyl (LHY), Circadian Clock Associated 1 (CCA1), Timing of CAB Expression 1 (TOC1)* and *Pseudo-Response Regulator (PRR1)*) exhibit conserved phasing and regulatory architecture comparable to those of temperate model species such as *Arabidopsis thaliana* (Zdepski et al., 2008). Time-of-day-specific cis-regulatory elements were also highly conserved, suggesting that circadian regulation has played a fundamental role in plant genome evolution irrespective of latitude or climate regime (Zdepski et al., 2008). Similarly, studies in tropical tree species have shown that key metabolic outputs, including volatile isoprenoid emissions, are under strong circadian control, highlighting the functional relevance of the clock in tropical forests (Wilkinson et al., 2006).

Despite these advances, our understanding of circadian clock behavior in tropical organisms remains highly fragmented. In particular, comparative analyses of circadian clock architecture across tropical and temperate lineages are largely lacking. It remains unclear whether core properties of the clock such as oscillation amplitude, phase stability, and robustness under fluctuating environmental conditions are conserved or systematically modified in tropical species, where seasonal variation in photoperiod and temperature is markedly reduced. Addressing this gap is essential for understanding how circadian clocks have evolved under different climatic regimes and for elucidating their ecological and evolutionary roles beyond temperate model systems.

*Rubroshorea leprosula* (previously *Shorea leprosula*), a dipterocarp widely distributed across Southeast Asian rainforests, is a representative model species for exploring circadian regulation in tropical trees. The humid tropical rainforests of Southeast Asia are characterized by a preponderance of trees of the Dipterocarpaceae family (Ashton, 2003). Dipterocarp trees are highly valued for both their contribution to forest diversity and their use in timber production and are well known for exhibiting a community-wide synchrony in flowering and fruiting (Ashton et al., 1988; Sakai et al., 1999; Chen et al., 2018; Numata et al., 2022). Their ecological importance, seasonal flowering patterns, and emerging genomic resources make it an ideal system to investigate how diurnal rhythms operate under low seasonal variation in day length and temperature.

In this study, we conducted a comprehensive time-series transcriptome analysis of *R. leprosula* to elucidate its circadian transcriptional landscape. Specifically, we (i) identified rhythmic genes using genome-wide expression profiling, (ii) classified rhythmic expression patterns through clustering, (iii) functionally characterized gene clusters via Gene Ontology enrichment, (iv) compared rhythmic dynamics of conserved clock genes between *Arabidopsis* and *R. leprosula*, and (v) reconstructed regulatory interaction networks to infer potential coordination among rhythmic genes. Together, these analyses provide the first transcriptomic framework of circadian regulation in a tropical tree, offering insights into how rhythmic expression and regulatory network organization may differ from temperate model species.

## 2. Materials and Methods

### 2.1 Sampling method, RNA extraction, and RNA-seq: 4-hour intervals over 48 hours

An individual of *R. leprosula* was germinated in a pot and cultivated in the nursery of Forest Research Institute Malaysia. The sapling pot was transferred to a light room at the University of Malaya, Malaysia, and grown at room temperature under a 12-hour light/12-hour dark cycle from September to November of 2019 (Fig. 1a). On November 12 at 7:00 pm (the timing of light-to-dark transition), the pot was placed in a growth chamber (GC-1600, Tech Lab Scientific Pty Ltd, Malaysia) covered with a black curtain and maintained at 25°C. The light was on only once, from 7:00 am to 7:00 pm on November 13, after which it remained in a constant dark environment (Fig. 1a).

**Figure 1.**
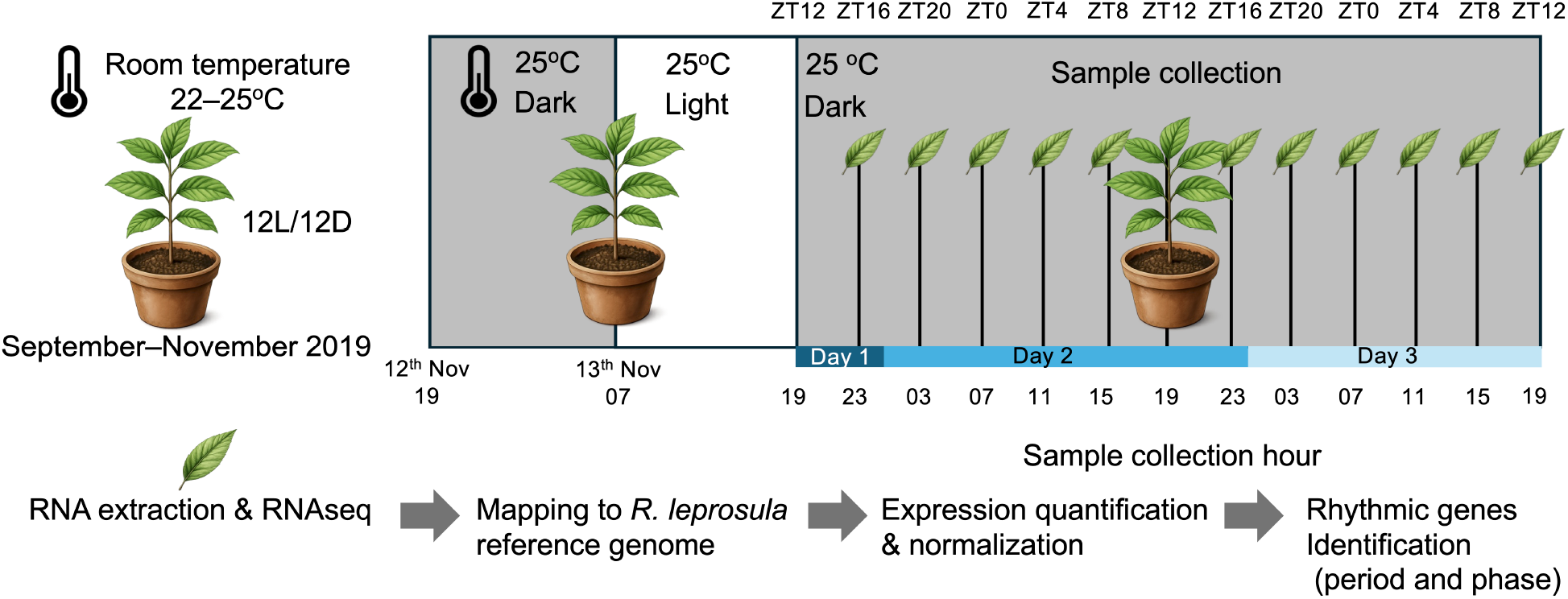
Identification of rhythmic genes in *R. leprosula*. Representative workflow for RNA-seq sampling, processing, and circadian rhythmicity analysis in *R. leprosula*.

Thirteen times of sampling were conducted between 7:00 pm on 13 November (immediately after the light was switched off) and 7:00 pm on 15 November, with an interval of 4 hours. In each sampling, three leaf slices were cut from an uninjured leaf with scissors under a dim green torch. Each slice was immediately put into a tube filled with RNAlater solution (Thermo Fisher Scientific) then stored in a - 80°C deep freezer. Total RNA was extracted from ca. 70 mg leaf samples in accordance with the CTAB method described in a previous study (Yeoh et al., 2017). RNA yield was determined using a NanoDrop ND-1000 (Thermo Fisher Science, USA). The extracted RNA was sent to Pacific Alliance Lab (Singapore), where a cDNA library was prepared with a NEBNext^®^ Ultra™ RNA Library Prep Kit for Illumina (New England BioLabs, Ipswich, MA, USA) and 150 paired-end transcriptome sequencing was conducted using an Illumina NovaSeq6000 sequencer (Illumina, San Diego, CA, USA).

### 2.2 RNA-seq Processing and Rhythmicity Detection

Raw sequencing data underwent a multi-step processing pipeline. File integrity was verified via MD5 checksum validation (check_md5.pl). Quality control and adapter trimming were performed using *fastp*, removing reads with >40% low-quality bases (Q<20) or length <50 bp (Chen et al., 2018). Clean reads were aligned to the *R. leprosula* reference genome using *STAR* (Dobin et al., 2013; Satake et al., 2024). The genome index was generated with --runMode genomeGenerate (overhang = 149 bp) using the annotated FASTA and GTF files. *STAR* was executed with --quantMode TranscriptomeSAM to produce transcriptome-aligned BAMs for downstream quantification.

Expression quantification was done using *RSEM* with the duplicate reference files to obtain expected counts, TPM, and FPKM values (Li & Dewey, 2011). Gene-level expected counts were imported into *edgeR* in R, and genes with mean RPK ≥ 5 were retained (Robinson et al., 2010). Normalisation was done via the TMM (Trimmed Mean of M-values) method, and log_2_-transformed CPM values were exported for rhythmicity analysis. Circadian rhythmicity was assessed using the RAIN (Rhythmicity Analysis Incorporating Nonparametrics) and JTK cycle algorithms in R (Thaben & Westermark, 2014; Hughes et al., 2010). The script defined 13 time points (0-48h at four hours intervals), executed *RAIN* genewise, and output period, phase, and adjusted p-values. Significant rhythmic genes were visualized using rose plots and expression heatmaps.

### 2.3 Rhythmic Gene Categorization and Heatmap Clustering

The subset of rhythmic genes identified by *RAIN* was extracted into a log_2_-CPM expression matrix and standardized by z-score transformation (mean = 0, SD = 1). Hierarchical agglomerative clustering was performed using Ward’s minimum variance method (method = “ward.D2”) with Euclidean distance. Based on the approximate expression phase distribution, four clusters (k = 4) were defined from the dendrogram. Heatmaps were generated with rows annotated by cluster membership and columns arranged by sampling time to preserve diurnal progression.

### 2.4 Ortholog Identification and GO Annotation Transfer

Putative orthologs between *R. leprosula* and *A. thaliana* were identified using *BLASTP* (E-value ≤ 1e-5). The best-scoring *Arabidopsis* hit for each *Rubroshorea* gene was retained to enable functional annotation transfer. Gene Ontology (GO) terms were assigned to *Rubroshorea* genes based on their *Arabidopsis* orthologs, focusing on the Biological Process (BP) category with evidence codes limited to Inferred from Electronic Annotation (IEA). The mapping of *Rubroshorea* genes to GO terms was converted into *GOstats*-compatible objects for enrichment analysis.

### 2.5 Gene Ontology Enrichment Analysis

Each rhythmic gene cluster (Section 2.3) served as a target set for enrichment analysis, while all annotated *R. leprosula* genes formed the background universe. Analysis was restricted to the Biological Process ontology. Over-representation was tested using Fisher’s exact test implemented in *GOstats* with conditional = TRUE to account for the GO hierarchy (Falcon & Gentleman, 2007). P-values were adjusted using the Benjamini-Hochberg method, and terms with adjusted p < 0.05 were deemed significantly enriched. For each cluster, enrichment outputs included odds ratios, observed and expected counts, and associated gene lists. The top five significantly enriched terms were visualized as bar plots (number of target genes per term) in R using *ggplot2*.

### 2.6 Phylogenetic and synteny analysis of clock genes

The alignment and phylogenetic tree construction were performed using the build function of ETE3 v3.1.3 (https://etetoolkit.org). In ETE3 build function, MAFFT v6.861b (https://mafft.cbrc.jp) was used for sequence alignment, trimAl v1.4.rev (https://vicfero.github.io/trimal) trimmed alignment gap, and phylogenetic tree construction was performed with RAxML v8.2.11 (https://cme.h-its.org/exelixis/web/software/raxml) using the GAMMA JTT model and 500 bootstrap replications. The resulting tree was visualized by iTOL (https://itol.embl.de).

Genomic synteny between *R. leprosula* and *P. chinensis* was investigated. First of all, we aligned all contig sequences of *R. leprosula* against *P. chinensis* using D-Genies (https://dgenies.toulouse.inra.fr/), and picked up contigs corresponding to chromosomes of *P. chinensis*. Next, we performed all-vs-all BLASTp search between all protein sequences on chromosomes for both species with a cut-off e-value of 1E-5. Then, MCScanX (https://github.com/wyp1125/MCScanX) was performed to detect syntenic blocks within or between genomes using the results of the above BLASTp searches. The collinear relationship between two species was shown by SynVisio (https://synvisio.github.io).

### 2.7 Comparative Analysis of Core Clock Genes

Twenty-one well characterized *A. thaliana* core clock genes (including *Late Elongated Hypocotyl (LHY), Circadian Clock Associated 1 (CCA1)*, members of the *Pseudo-Response Regulator (PRR)* family including *PRR9, PRR7, PRR5*, and *Timing of CAB Expression 1 (TOC1/PRR1), Early Flowering 4 (ELF4), Lux Arrhythmo (LUX), GIGANTEA (GI), ZEITLUPE (ZTL), REVEILLE (RVE)* transcription factors *(RVE8, RVE6*, and *RVE4*), among others) were selected to represent morning, afternoon, and evening modules. Their putative *R. leprosula* orthologs were identified using *BLASTP* searches against the translated *Rubroshorea* proteome. Expression profiles of *Arabidopsis* orthologs under constant dark conditions were obtained from the *DIURNAL* database (Mockler et al., 2007).

Comparative expression analyses evaluated rhythmic parameters in both species using (i) relative amplitude (amplitude normalised to mean expression) and (ii) coefficient of variation (CV) across time points, alongside phase and period estimates. These metrics enabled quantitative assessment of circadian conservation or divergence between *A. thaliana* and *R. leprosula*.

To further interpret rhythmic coordination at the systems level, we reconstructed the *Rubroshorea* gene-interaction network using time-series correlations.

### 2.8 Correlational network extraction

A directed gene-interaction network was inferred from *R. leprosula* time-series expression (13 time points, four-hour intervals) using time-lagged cross-correlation (TLCC). Missing values were linearly interpolated, and each gene’s expression was z-scored across time. Cross-correlations were computed for every ordered gene pair for lags from −6 to +6 (−24 to +24 h). Directionality was assigned based on the lag yielding the highest absolute correlation (positive lag: source → target).

Edges were retained if they met the following criteria: |lag| ≥ 1, |correlation| ≥ 0.55, |zero-lag correlation| ≤ 0.35, and margin over second-best lag ≥ 0.05. Edge weights were set to the absolute correlation coefficient, and networks were optionally filtered to the top 500 edges. The resulting directed network was visualised using a hierarchical layout in *Cytoscape* (v3.10). Node level metrics were computed to identify potential regulatory hubs, including degree, strength, PageRank, and betweenness centrality.

## 3. Results

### 3.1 Rhythmic Genes Identified and Period Distribution

To investigate circadian transcriptional regulation in *R. leprosula*, we first profiled genome-wide rhythmicity using the RAIN algorithm. Out of 20,748 expressed genes, 283 exhibited significant rhythmic oscillations under constant dark conditions. This represents ∼1.3% of the transcriptome, which is slightly less than to ∼1.6% of transcriptome in Arabidopsis under constant dark condition, suggesting only a subset of genes is under strong circadian control (Wenden et al., 2011). A heatmap of 20,748 expressed genes highlights broad expression dynamics but does not reveal distinct rhythmic oscillations in most genes (Supplementary figure 1).

The estimated circadian periods of rhythmic genes spanned 21-27 h, with the majority clustering between 23 and 25 h (Figure 2a). Genes with shorter (<23 h) or longer (>25 h) periods were comparatively underrepresented. This distribution indicates that the *Rubroshorea* circadian system predominantly maintains ∼24 h periodicity, consistent with canonical circadian rhythms, but also allows flexibility across a broader temporal range.

**Figure 2.**
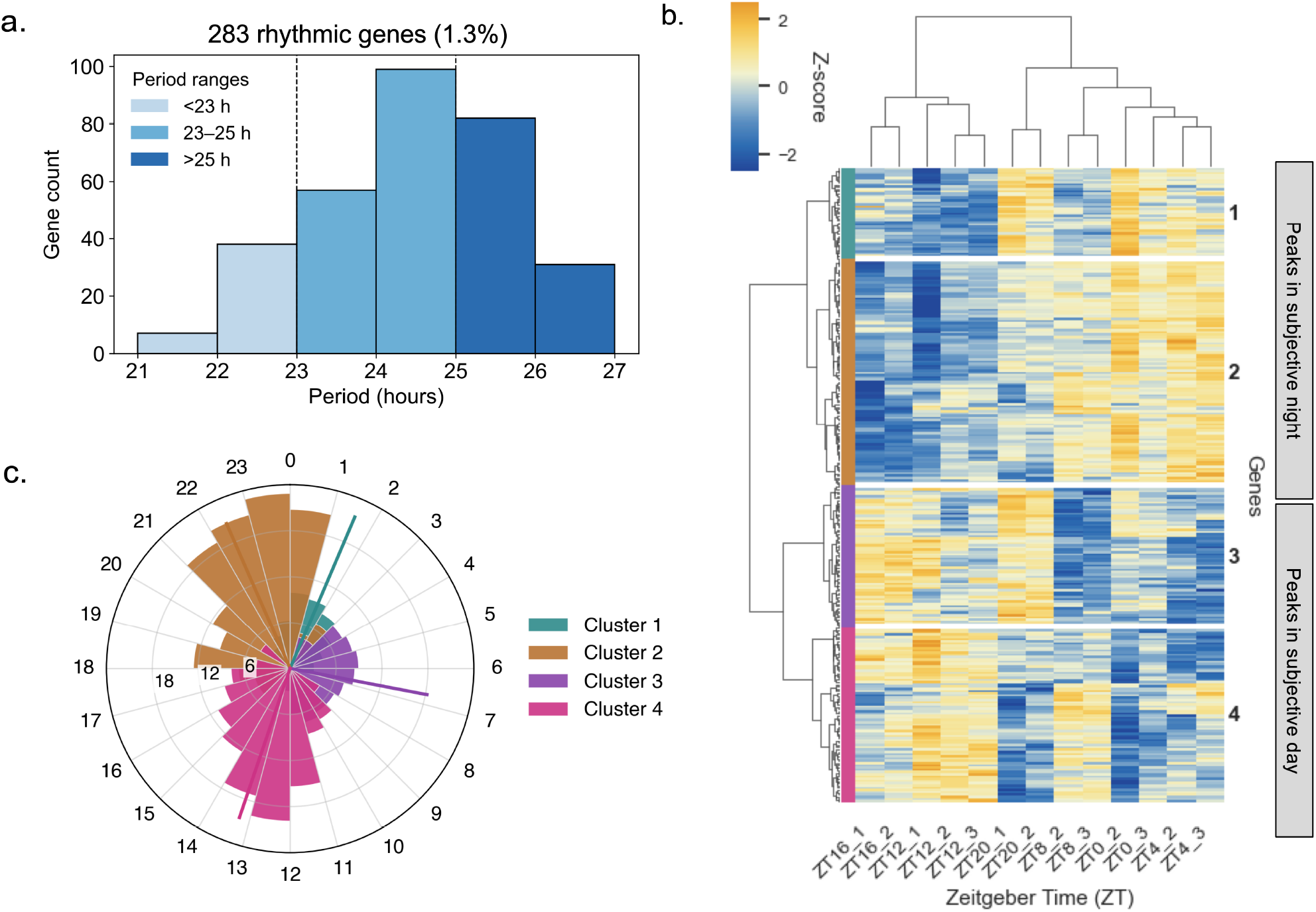
Clustering and functional analysis of rhythmic genes. (a) Distribution of circadian periods among rhythmic genes, grouped into <23 h, 23-25 h, and >25 h categories. (b)Heatmap of 283 rhythmic genes across Zeitgeber time (ZT), organized into four major clusters (suffix number represents number of passed days). (c) Combined rose plot showing the phase distribution of rhythmic clusters, color-coded by cluster.

### 3.2 Clustering and functional analysis of rhythmic genes

Hierarchical clustering of the 283 rhythmic genes revealed four major expression clusters (Figure 2b). Each cluster displayed distinct phase-enriched expression peaks across the diurnal cycle, reflecting different temporal modules of circadian regulation. Phase analysis using rose plots highlighted clear separation between clusters (Figure 2c). Cluster 1 genes peaked around early subjective morning, Cluster 2 in the late morning to midday, Cluster 3 in the afternoon, and Cluster 4 during the night. This phase-resolved organization indicates that rhythmic genes in *R. leprosula* are temporally partitioned, similar to the phased waveforms described in other plant species.

GO enrichment analysis revealed that rhythmic gene clusters are functionally distinct (Figure 3). Cluster 1 was enriched for light- and chloroplast-related processes such as *photomorphogenesis* and *chloroplast rRNA processing*. Cluster 2 showed enrichment for *response to blue light, ethylene biosynthesis*, and *vascular transport*, indicating links to light signaling and hormone regulation. Cluster 3 was associated with *mitochondrial genome maintenance, peptide biosynthesis*, and *cytokinin response*, highlighting roles in energy metabolism and hormone sensing. Cluster 4 was enriched for protein quality control and organelle-associated processes, including *mitotic spindle disassembly, mitochondrial protein processing*, and *autophagosome maturation*.

**Figure 3.**
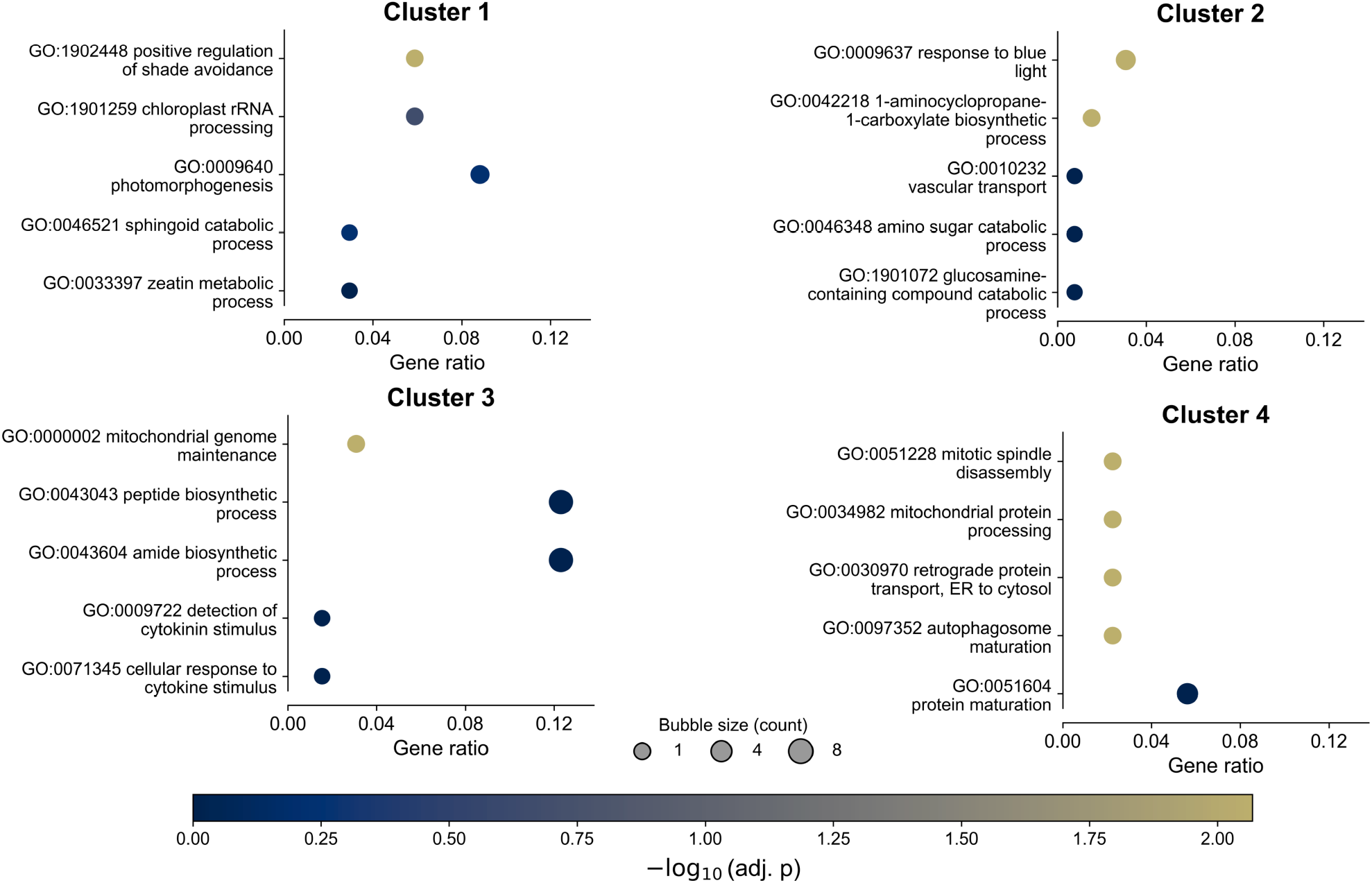
Bubble plots of enriched Gene Ontology Biological Process (GOBP) terms for each cluster. Bubble size indicates gene count, and colour represents statistical significance (-log10 adjusted *p* value). Distinct processes are enriched in each cluster, including light and chloroplast functions (Cluster 1), blue light and hormone-related processes (Cluster 2), mitochondrial and cytokinin-related processes (Cluster 3), and protein processing and quality control (Cluster 4).

These results show that rhythmic genes in *R. leprosula* are organized into distinct temporal clusters that not only differ in their phase of expression but also in their functional specializations, ranging from light signaling to metabolism and cellular homeostasis.

### 3.3 Identification and Comparative Analysis of Circadian Clock Genes

To identify candidate circadian regulators in *R. leprosula*, we used Arabidopsis core clock components as queries and retrieved putative *Rubroshorea* orthologs by sequence homology. A total of 21 *Arabidopsis* clock genes, including *LHY, CCA1, PRR9, PRR7, PRR5, TOC1, ELF4, LUX, GI, ZTL, RVE8, RVE6*, and *RVE4*, were matched to multiple *Rubroshorea* orthologs (Figure 4a; Supplementary Table 1). Several genes showed more than one *Rubroshorea* copy, consistent with gene duplication events in the species.

**Figure 4.**
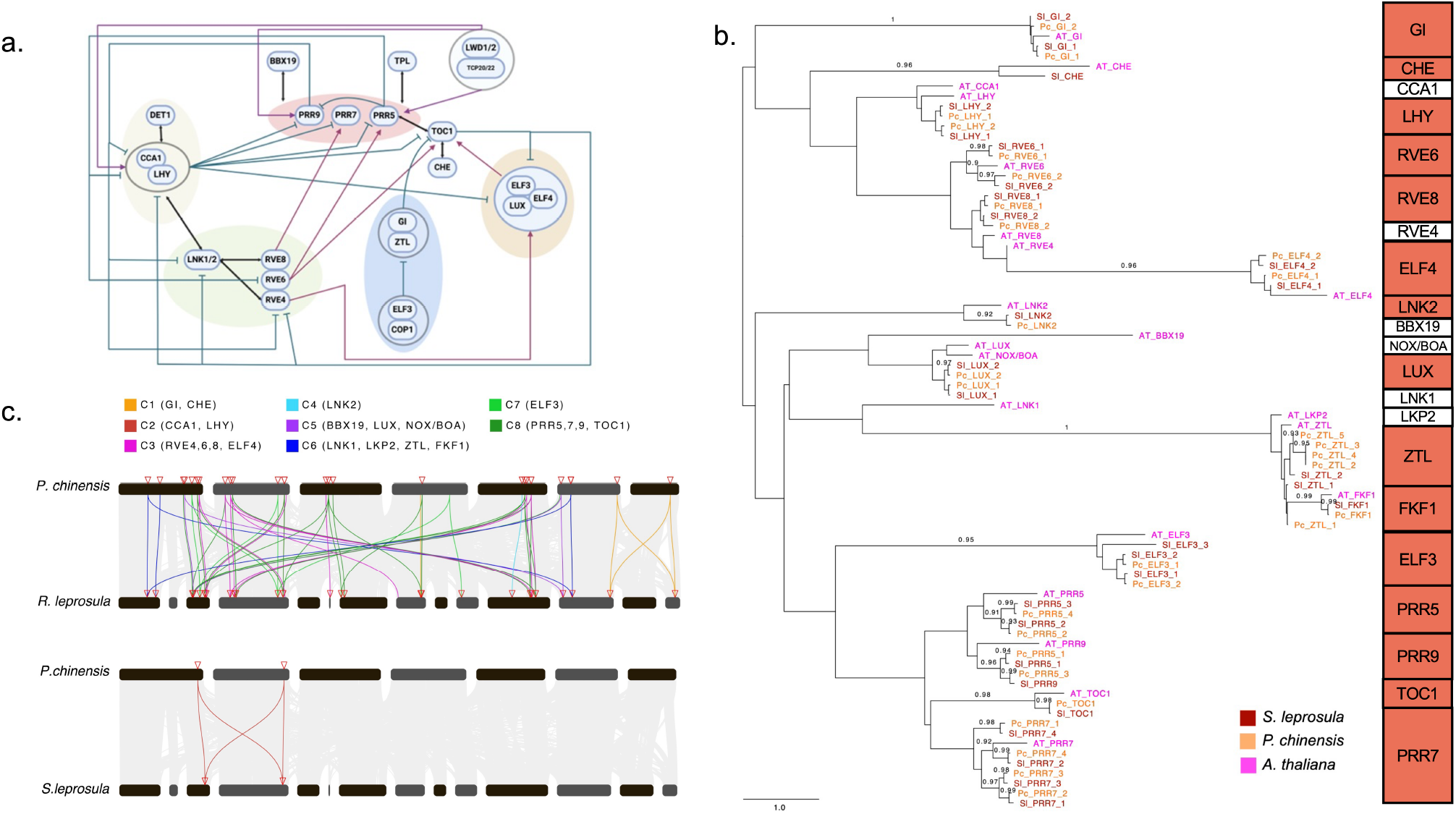
Comparative analysis of circadian clock genes in *R. leprosula*. (a) Schematic overview of the *Arabidopsis* circadian clock network, highlighting core morning, day, and evening components and associated regulatory interactions (Singh & Srivastava, 2024). (b) Phylogenetic analysis confirms the placement of *Rubroshorea* orthologs within well-supported clades alongside *A. thaliana* and *P. chinensis*. (c) Conserved syntenic blocks between *P. chinensis* and *R. leprosula* containing core circadian clock genes. Coloured connections denote collinear orthologs (C1-C8), and red triangles indicate clock gene positions. Preservation of gene order around *LHY* and *PRR* loci indicates retention of ancestral circadian genomic organization in Dipterocarpaceae.

Phylogenetic analysis placed *Rubroshorea* orthologs alongside their *Arabidopsis* and *Parashorea chinensis* counterparts in well-supported clades (Figure 4b). For example, *Rubroshorea LHY* grouped tightly with *Arabidopsis* and *P. chinensis LHY* sequences that is a close relative of *R. leprosula*, while multiple *Rubroshorea PRR* genes clustered with the *PRR7* and *PRR9* subgroups, reflecting lineage-specific expansions. This analysis confirmed that *Rubroshorea* retains recognizable orthologs of major clock gene families.

To further investigate conservation, we performed synteny analysis between *Rubroshorea* and *P. chinensis*. Several clock genes, including *LHY* and *PRR* members, were located within conserved syntenic blocks between the two species (Figure 4c). This highlights both structural conservation and possible retention of ancestral circadian loci within the Dipterocarpaceae lineage.

Together, these analyses demonstrate that *R. leprosula* possesses clear orthologs of *Arabidopsis* clock genes, with evidence of duplication and conservation at both phylogenetic and structural levels. This provides a solid foundation for exploring circadian regulation in this tropical tree species.

### 3.4 Expression Dynamics and Network Inference of *Rubroshorea* Clock Genes

We first examined the temporal expression profiles of core clock genes in *R. leprosula* alongside their *A. thaliana* orthologs (Figure 5a). Core components, including *LHY, GI, LUX, PRR5*, and *PRR7*, displayed clear circadian oscillations in *Rubroshorea*, with peak phases broadly aligned with those observed in *Arabidopsis*. To quantify rhythmic strength, oscillation amplitude was defined as the peak-to-trough difference of normalized expression levels across the diurnal cycle. Using this metric, *Rubroshorea* orthologs consistently exhibited reduced amplitudes relative to their *Arabidopsis* counterparts, together with smoother temporal profiles, indicating attenuated transcriptional oscillations characteristic of this tropical species.

**Figure 5.**
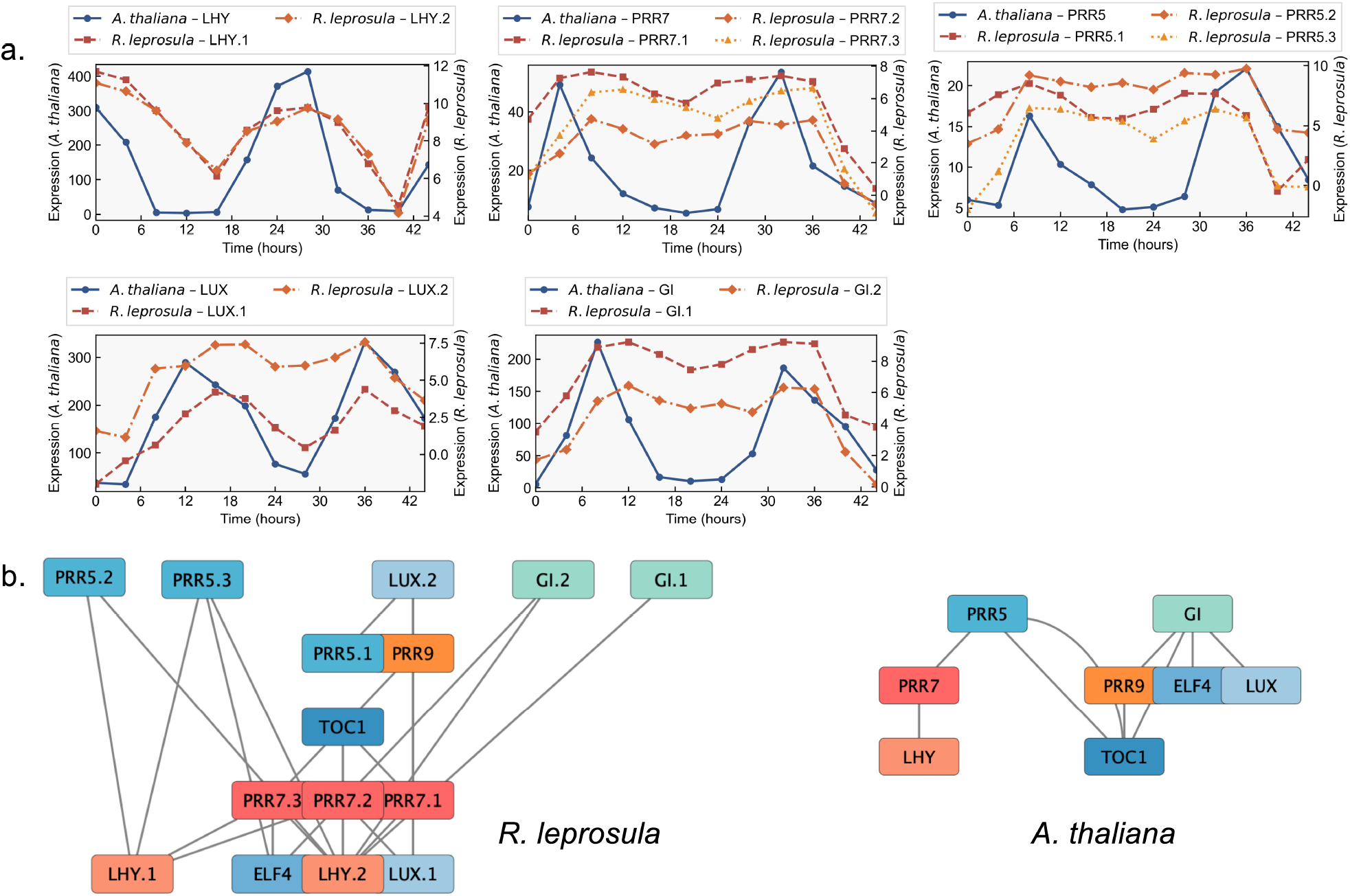
Expression profiles and network inference of clock genes. (a) Temporal expression profiles of key clock genes (*LHY, GI, LUX, PRR5*, and *PRR7*) in *R. leprosula* compared with Arabidopsis under diurnal conditions. While phases are conserved, *Rubroshorea* orthologs display reduced amplitudes. (b) Networks of clock gene interactions reconstructed using the time-lagged cross-correlation (TLCC) method under constant dark conditions. Networks are arranged using hierarchical clustering for both *Rubroshorea* (left) and *Arabidopsis* (right), showing overall similarity in organization despite the presence of paralogs in *Rubroshorea*.

To investigate regulatory architecture, we applied time-lagged cross-correlation (TLCC) to gene expression profiles under constant dark conditions and reconstructed directed interaction networks for *R. leprosula* and *A. thaliana* (Figure 5b). Network similarity was assessed quantitatively by comparing shared directed edges, node connectivity patterns, and hierarchical clustering structure derived from TLCC peak correlations and associated time lags. Using these metrics, both species exhibited a high degree of overlap in core regulatory interactions, including conserved links among *LHY, PRR* family members, *GI, LUX*, and *ELF4*. Despite the presence of multiple paralogs in *Rubroshorea* (e.g., *PRR5, PRR7*, and *LHY*), which increased network size and redundancy, paralogous nodes clustered with their corresponding functional groups, preserving the canonical circadian architecture observed in *Arabidopsis*.

These results demonstrate that despite differences in rhythmic amplitude, the underlying network topology of circadian genes in *R. leprosula* remains highly conserved with Arabidopsis, supporting evolutionary conservation of the core circadian oscillator.

### 3.5 Comparative Robustness of Clock Gene Expression

To assess differences in the strength and stability of circadian rhythms, we compared the temporal expression profiles of core clock orthologs between *A. thaliana* and *R. leprosula*. Two metrics were evaluated: the coefficient of variation (CV), which reflects variability in rhythmic expression, and relative amplitude, which measures the strength of oscillation relative to baseline expression.

*Rubroshorea* orthologs showed consistently lower CV values than Arabidopsis genes (Figure 6; *p* = 0.022, using the Mann-Whitney U test), indicating reduced variability across the circadian cycle. Likewise, relative amplitudes were significantly lower in *Rubroshorea* (Figure 6; *p* = 0.037, using the Mann-Whitney U test), suggesting that oscillations are less pronounced in this tropical tree species. These patterns point to a general dampening of transcriptional rhythms in *Rubroshorea* compared to *Arabidopsis*.

**Figure 6.**
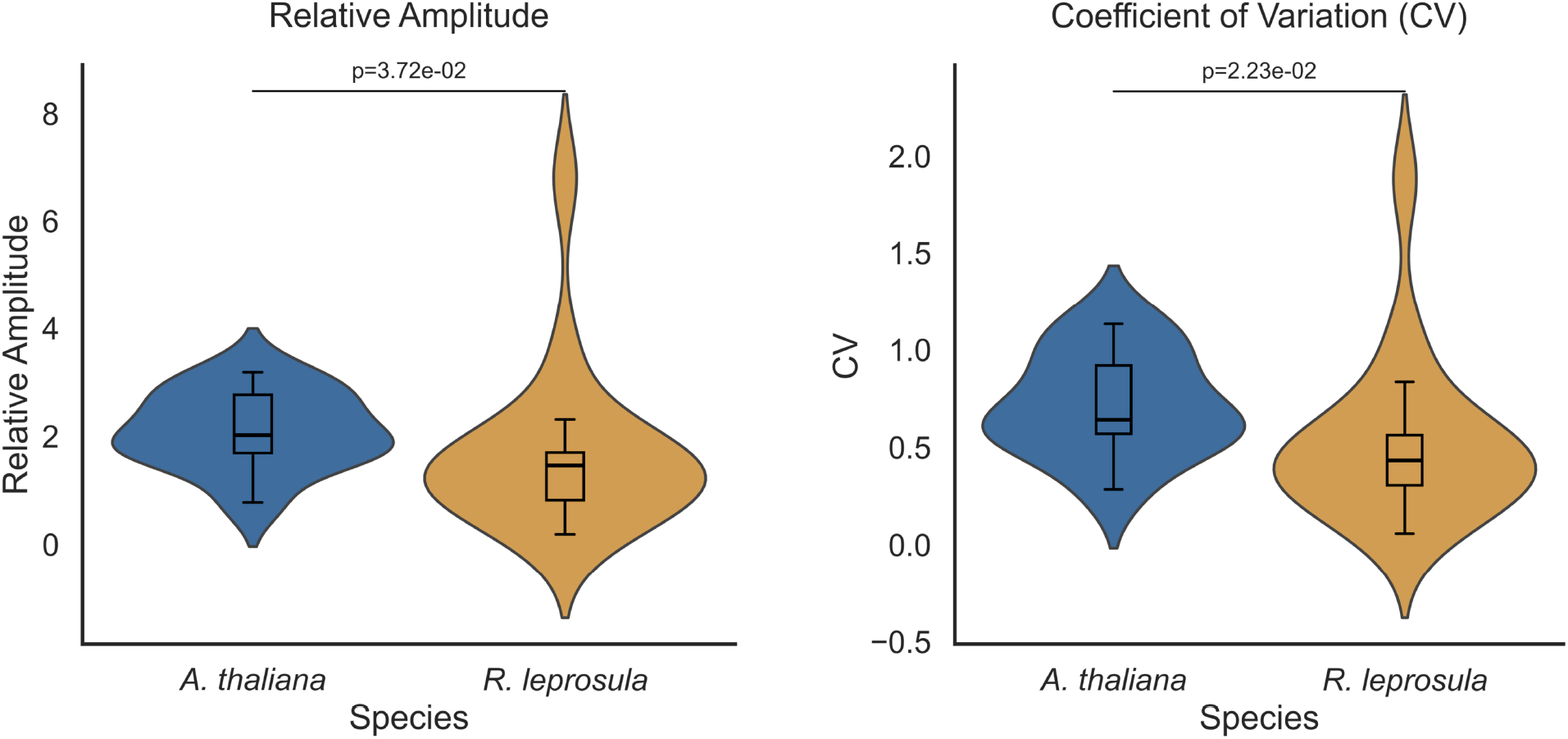
Comparative robustness of circadian expression between species. Boxplots comparing (left) coefficient of variation (CV) and (right) relative amplitude of core clock gene expression in Arabidopsis and *R. leprosula. Rubroshorea* shows significantly lower variability (*p* = 0.0223) and lower amplitude (*p* = 0.0372), indicating dampened transcriptional rhythms relative to Arabidopsis.

Together with the expression profiles and network analysis, these results suggest that while *R. leprosula* maintains a conserved circadian framework, its rhythmic outputs are more buffered and less robust. This may reflect species-specific adaptations, where weaker oscillations are sufficient for circadian regulation in the relatively stable tropical environment, in contrast to the stronger rhythms observed in Arabidopsis under temperate conditions.

## 4. Discussion

In this study, we characterized the circadian transcriptional landscape of *R. leprosula*, a tropical dipterocarp tree, and compared it with the well-established circadian framework of *A. thaliana*. Our genome-wide rhythmicity analysis identified a distinct subset of genes (∼1.3% of the transcriptome) exhibiting robust circadian oscillations, indicating that clock-controlled regulation in *R. leprosula* is selective rather than globally pervasive. This proportion is substantially lower than that reported for herbaceous species such as *Oryza sativa* (rice), where approximately 22% of expressed genes show rhythmic behavior under similar conditions (Zhang et al., 2023). Rather than reflecting weaker day and night entrainment, this reduced rhythmic output in *Rubroshorea* likely arises from long-term adaptation to tropical environments, where daily light and dark cycles remain strong but seasonal variation in photoperiod and temperature is minimal. Under such conditions, circadian regulation may be concentrated on a core set of functionally essential genes, while broader transcriptomic rhythms often associated with seasonal anticipation and developmental plasticity are less prominent.

The estimated period distribution, centered around ∼24 h, supports the existence of an endogenous circadian rhythm that aligns with the canonical circadian cycle. The narrow clustering of periods between 23-25 h, along with fewer genes exhibiting shorter or longer oscillations, suggests that *Rubroshorea* maintains a well-tuned internal clock with limited period variability. Such constrained rhythmicity may facilitate stable timekeeping in the tropical environment, where environmental oscillations are less variable than in temperate regions.

Hierarchical clustering revealed four distinct temporal modules of gene expression, each enriched for specific biological processes. Early-day clusters were dominated by genes involved in light signaling and chloroplast function, whereas midday and afternoon clusters included genes related to hormone response, mitochondrial activity, and peptide biosynthesis. The night-phase cluster was associated with protein quality control and autophagy-related processes. This temporal partitioning highlights how *Rubroshorea* organizes physiological processes across the day-night cycle, enabling coordinated regulation of photosynthesis, metabolism, and cellular maintenance. The clear phase separation of rhythmic clusters mirrors patterns observed in Arabidopsis and other model plants, underscoring the conserved modular organization of the plant circadian transcriptome.

Comparative analysis of circadian clock genes further demonstrated strong evolutionary conservation across *Rubroshorea, Arabidopsis*, and *Parashorea chinensis*. Phylogenetic and synteny analyses confirmed that Shorea retains orthologs of all major clock gene families including LHY, CCA1, PRR, TOC1, ELF4, LUX, GI, and ZTL with evidence of lineage-specific duplications. The high similarity in exon and intron structures with syntenic relationships suggests that these regulators have maintained their functional roles since their divergence from common ancestors within Dipterocarpaceae. Despite this structural and phylogenetic conservation, rhythmic expression in Shorea was notably dampened compared to Arabidopsis. Lower amplitude and smoother oscillations were observed in *Rubroshorea* orthologs, particularly for key morning and evening genes. The reduced coefficient of variation and relative amplitude point toward weaker rhythmic robustness, possibly reflecting adaptive attenuation of circadian strength in response to the stable tropical environment. A similar reduction in amplitude has been reported in other tropical or shade-adapted species, suggesting that strong circadian oscillations may not be required for survival under minimal photoperiodic variation.

Network inference using the time-lagged cross-correlation (TLCC) approach revealed that Shorea retains a canonical circadian architecture comparable to Arabidopsis. The core regulatory links among LHY, CCA1, PRR family members, GI, LUX, and ELF4 were preserved, indicating conservation of interlocking feedback loops central to clock operation. Interestingly, the presence of multiple paralogs in Shorea expanded the network complexity without altering its overall topology, hinting at possible sub functionalization or redundancy among duplicated components. This conservation of topology, despite expression dampening, suggests that while the circadian machinery remains intact, its downstream rhythmic output may be modulated to suit Shorea’s ecological niche.

Overall, our results reveal that *R. leprosula* possesses a structurally conserved but functionally attenuated circadian system. Its core clock gene network mirrors that of Arabidopsis, yet with weaker rhythmic amplitude and reduced transcriptomic rhythmicity, reflecting a shift toward environmental-stability-driven circadian adaptation. This attenuation aligns with theoretical predictions from Seki and Ito (2022), who demonstrated that robust self-sustained circadian oscillators are selectively favoured in strongly seasonal environments, where large shifts in photoperiod necessitate an internal rhythm that can maintain phase independently of day length. In contrast, aseasonal tropical environments, such as those inhabited by R. leprosula, provide minimal selective pressure for maintaining high-amplitude, self-sustained oscillations. Under such conditions, damped or externally reinforced rhythms are sufficient to ensure proper alignment with largely invariant dawn-dusk cycles. The transcriptomic features we observe-low rhythmic gene proportion, narrow period distribution, and reduced amplitude-are consistent with this evolutionary framework, suggesting that Shorea’s circadian system has adapted toward stability, energetic efficiency, and reliance on reliable environmental cues rather than on a dominant endogenous oscillator. This interpretation is consistent with observations from other low-latitude systems. For example, duckweeds exhibit increasingly irregular circadian behavior toward equatorial regions (Isoda et al., 2022), while some cyanobacteria have lost circadian rhythmicity entirely in stable tropical environments (Holtzendorff et al., 2008; Flombaum et al., 2013). These findings support the idea that reduced environmental seasonality can relax selective pressure for maintaining strong endogenous oscillators, aligning with the attenuated rhythmic transcription observed in *R. leprosula*. These findings provide new insight into how tropical trees balance internal rhythmic regulation with the environmental predictability of equatorial forests, contrasting sharply with the photoperiod-tracking strategies favoured in temperate species.

## 5. Conclusion

This study presents the first comprehensive circadian transcriptional profile of *R. leprosula*, a dominant tropical dipterocarp species. By integrating genome-wide rhythmicity analysis, gene clustering, and comparative network inference, we reveal that *Rubroshorea* maintains a conserved circadian framework similar to that of *A. thaliana* yet exhibits distinct functional and rhythmic characteristics. The presence of clearly identifiable orthologs for all major clock components, together with conserved exon-intron structures and syntenic relationships, underscores the deep evolutionary conservation of the core oscillator.

Despite this conservation, *Rubroshorea* displays attenuated oscillatory strength and lower rhythmic variability, indicating a more buffered circadian system. Such dampened rhythms may represent an adaptive strategy to the stable light and temperature regimes of tropical forests, where extreme diel fluctuations are minimal. The temporal clustering of rhythmic genes into distinct functional modules from light and chloroplast processes to metabolic and protein homeostasis pathways illustrates how *Rubroshorea* coordinates physiological processes over the daily cycle while maintaining flexibility under relatively constant environmental conditions.

Importantly, this work provides a baseline framework for circadian biology in tropical trees, offering insights that extend beyond fundamental chronobiology. The identification of conserved but fine-tuned circadian regulation in *R. leprosula* has broader ecological and forestry implications, as circadian rhythms influence key traits such as growth rate, photosynthetic efficiency, and stress resilience. Understanding how tropical trees modulate their circadian outputs can inform strategies for forest management, reforestation, and adaptation to climate change, particularly in species-rich equatorial ecosystems where environmental cues are subtle yet ecologically significant.

In summary, our findings demonstrate that *R. leprosula* harbors a conserved yet modulated circadian network, reflecting evolutionary adaptation to tropical stability. This study not only establishes the molecular foundation for exploring circadian regulation in dipterocarps but also opens new avenues for linking circadian dynamics with ecological function and forest productivity in tropical ecosystems.

## 6. Limitations and Future Directions

While this study establishes a foundational understanding of circadian regulation in *R. leprosula*, several limitations remain that warrant further investigation. First, our analyses were based solely on transcriptomic data under a single environmental condition. Although rhythmicity detection identified clear oscillations, the absence of corresponding protein-level and physiological measurements limits direct inference of post-transcriptional or metabolic rhythmic control. Future studies integrating proteomic, metabolomic, and phenotypic datasets will be critical to fully capture circadian outputs and downstream regulatory effects.

Second, the current sampling design reflects a controlled diurnal condition and therefore may not capture the plasticity of the clock under variable photoperiods or temperature regimes. Experimental validation under different light-dark cycles, temperature fluctuations, or stress conditions will help reveal how the *Rubroshorea* clock responds to environmental perturbations typical of tropical ecosystems.

Third, although the time-lagged cross-correlation (TLCC) method provided valuable insight into gene-gene associations, it infers statistical rather than causal relationships. Incorporating perturbation-based approaches, dynamic modelling, or network inference using multi-omics time series would enhance understanding of directionality and feedback strength within the circadian system.

Finally, the comparative analysis was restricted to *A. thaliana* and *P. chinensis*. Expanding comparative studies to additional Dipterocarpaceae members or other tropical tree lineages could help distinguish lineage-specific adaptations from general tropical circadian features. Such phylogenetically broad investigations will improve understanding of how circadian regulation evolves in perennial woody plants.

Looking ahead, **f**uture research should focus on functional validation of key clock orthologs, such as *LHY, CCA1, PRRs*, and *ELF4/LUX*, using gene editing or overexpression in model systems or in *Rubroshorea* itself. Establishing transformation protocols and reporter assays will enable direct testing of oscillator function. Furthermore, linking circadian gene regulation with ecological and forestry traits including growth rhythms, flowering time, and stress tolerance will provide practical insights for tropical forest management and tree breeding.

By addressing these directions, future work can move beyond transcriptional characterization toward a mechanistic and ecological understanding of circadian adaptation in tropical trees, bridging molecular chronobiology with long-term forest resilience and productivity.

## Acknowledgement

This study was supported by JST Sakura Science Program awarded to S.K.S. and JSPS KAKENHI (JP21H04781, JP23H04965, and JP23H04966) awarded to A.S.

