## Supplementary File for "Circadian Timekeeping in the Tropics: Rhythmic Transcriptome and Diurnal Regulatory Networks in *Rubroshorea leprosula*"

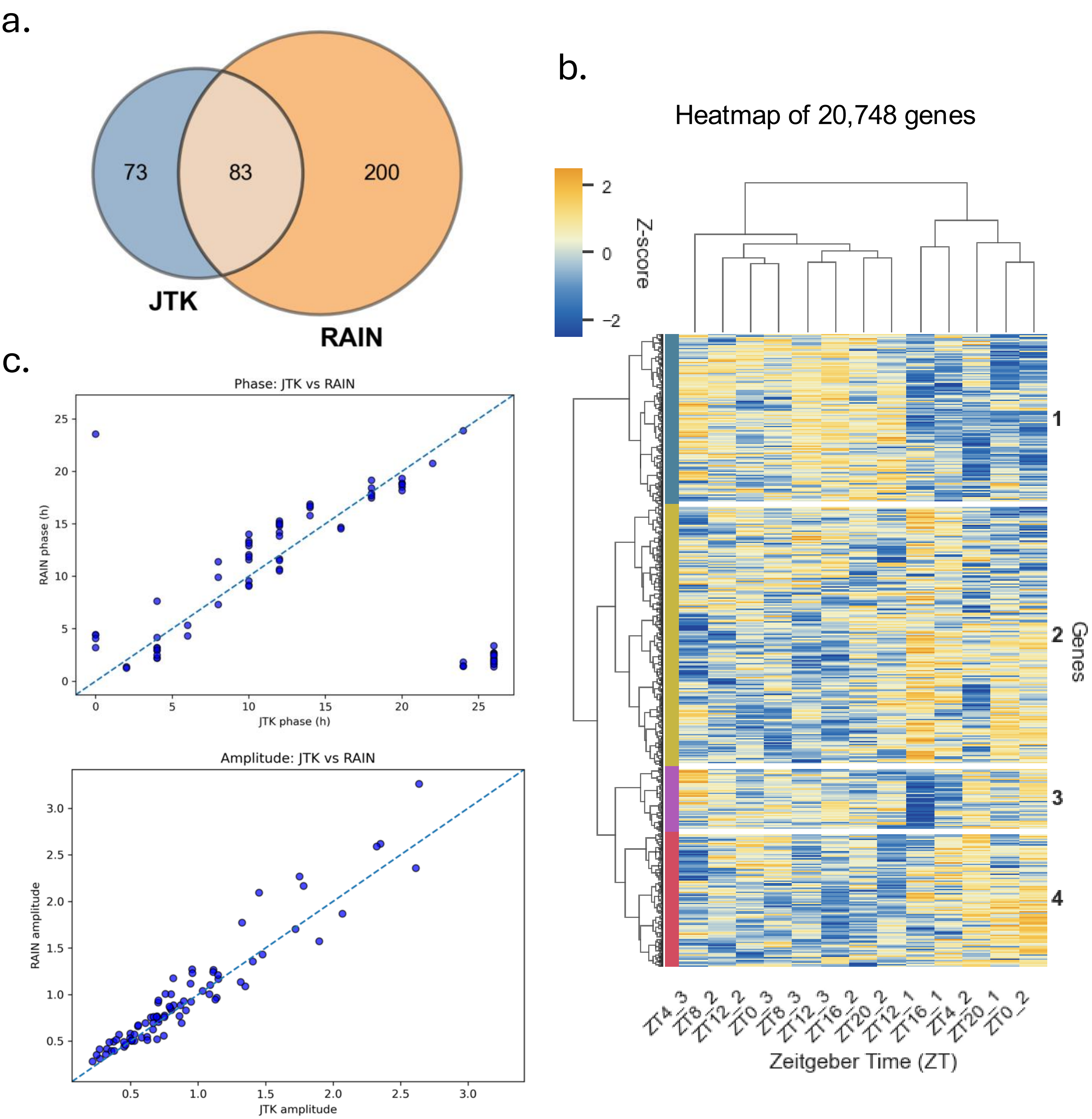


**Supplementary Figure 1 - Rhythmic gene identification and global expression patterns in Shorea leprosula**

**(a)** Overlap of rhythmic genes identified by JTK_CYCLE (JTK) and RAIN. A total of 83 genes were common, while 73 and 200 genes were uniquely detected by JTK and RAIN, respectively.

**(b)** Hierarchical clustering heatmap of all 20,748 genes across Zeitgeber time (ZT). Expression values are Z-score normalized, with genes grouped into four clusters based on temporal patterns.

**(c)** Comparison of phase (top) and amplitude (bottom) estimates between JTK and RAIN for commonly identified genes, showing strong agreement along the diagonal.

| Arath_gene | gene_id |
|---|---|
| LHY | AT1G01060 |
| PRR9 | AT2G46790 |
| PRR7 | AT5G02810 |
| PRR5 | AT5G24470 |
| TOC1 | AT5G61380 |
| LNK2 | AT3G54500 |
| FKF1 | AT1G68050 |
| ELF4 | AT2G40080 |
| LUX | AT3G46640 |
| GI | AT1G22770 |
| ZTL | AT5G57360 |
| RVE8 | AT3G09600 |
| CCA1 | AT2G46830 |
| BBX19 | AT3G21150 |
| CHE | AT5G08330 |
| NOX/BOA | AT3G46661 |
| RVE4 | AT5G02840 |
| RVE6 | AT5G52660 |
| ELF3 | AT2G25930 |
| LKP2 | AT2G18915 |
| LNK1 | AT2G43140 |

| Arath_ID | Gene name | Shorea_orthologs |
|---|---|---|
| AT1G01060 | *Homeodomain-like superfamily protein* | g3091, g18316 |
| AT1G22770 | *gigantea protein (GI)* | g33055, g24033 |
| AT1G68050 | *flavin-binding, kelch repeat, f-box 1* | g14398 |
| AT2G25930 | *hydroxyproline-rich glycoprotein family protein* | g3711, g18880, g38593 |
| AT2G40080 | *EARLY FLOWERING-like protein (DUF1313)* | g10832, g7216 |
| AT2G46790 | *pseudo-response regulator 9* | g3083 |
| AT3G09600 | *Homeodomain-like superfamily protein* | g7699, g10326 |
| AT3G46640 | *Homeodomain-like superfamily protein* | g19665, g28428 |
| AT3G54500 | *agglutinin-like protein* | g12409 |
| AT5G02810 | *pseudo-response regulator 7* | g10331, g11028, g6971, g19783 |
| AT5G08330 | *TCP family transcription factor* | g37582 |
| AT5G24470 | *two-component response regulator-like protein* | g18308, g28932, g37640 |
| AT5G52660 | *Homeodomain-like superfamily protein* | g18140, g40428 |
| AT5G57360 | *Galactose oxidase/kelch repeat superfamily protein* | g15121, g27345 |
| AT5G61380 | *CCT motif -containing response regulator protein* | g29220 |

**Supplementary Table 1 - Identification of Shorea leprosula orthologs of Arabidopsis circadian clock genes**

List of core and associated *Arabidopsis thaliana* circadian clock genes with their corresponding AGI identifiers and annotated gene names, alongside the identified *Shorea leprosula* orthologs. Orthologs were determined based on sequence similarity and mapping analyses. Multiple Shorea gene IDs are reported where applicable, reflecting potential gene duplication or paralogous relationships within the Shorea genome.
